# *In silico* Computed Clusters of Prostate Luminal Progenitors Match FACS-Enriched LSC^med^ cells

**DOI:** 10.1101/2021.06.16.448624

**Authors:** Emilia Puig Lombardi, Manon Baures, Charles Dariane, Jacques-Emmanuel Guidotti, Vincent Goffin

## Abstract

Several groups recently published single-cell (sc) expression atlases of the adult mouse prostate cells based on RNA sequencing (scRNA-seq) data. All studies identified one computerized cluster of non-secretory luminal progenitor cells enriched in luminal and stemness-related gene transcripts. The actual correspondence between these luminal progenitor cell clusters has not been investigated. In addition, the presence of *Krt4* (encoding cytokeratin 4) in these *in silico*-identified luminal progenitors suggested the overlap with FACS-enriched LSC^med^ luminal progenitor cells earlier identified as a stem-like, castration-tolerant and tumor-initiating cell population. Here, we used a unified bioinformatics pipeline to re-analyze published prostate scRNA-seq datasets and perform various pan-transcriptomic comparisons including the LSC^med^ cell signature. Our study demonstrates that i) the mouse prostate luminal progenitor cell clusters identified in the different scRNA-seq studies largely overlap and can be defined by a common 15-gene signature including *Krt4*, ii) mouse LSC^med^ cells match both mouse and human luminal progenitors identified by scRNA-seq analysis. Bridging these *in silico-*identified and *ex vivo-*characterized prostate luminal progenitor subsets should benefit our understanding of their actual involvement in prostate diseases.

## Introduction

Prostate diseases include benign (prostatitis, benign prostate hyperplasia) and malignant (prostate cancer) pathologies. The cells-of-origin of these pathologies are still poorly understood and their identification is the object of intense investigations to improve treatments. The prostate gland contains a glandular compartment surrounded by a stroma. The glandular epithelium contains two main cell types named basal and luminal according to their position in the pseudostratified epithelium, and very rare neuroendocrine cells (Abate-Shen and Shen 2000). Adult prostate stem cells have been hypothesized to mainly reside within the basal cell layer (English et al. 1987, Evans and Chandler 1987, Lawson et al. 2007). Subsequent studies suggested that rare luminal cells could also display stemness and initiate prostate tumors (Wang et al. 2009, Wang et al. 2014). In 2014, we discovered and isolated by cell sorting a rare population of luminal cells exhibiting stem/progenitor properties. We named these cells LSC^med^ according to their FACS profile (<u>L</u>in^-^/<u>S</u>ca-1^+^/<u>C</u>D49f<u>^med^</u>) using stem cell antigen-1 (Sca-1) and CD49f (integrin α6) as cell surface markers (Sackmann-Sala et al. 2014). LSC^med^ cells express typical luminal genes (e.g. cytokeratin (CK) 8 and androgen receptor) and exhibit stem/progenitor-like properties being able to generate spheres and organoids *in vitro*, and glandular structures when engrafted into host mice (Sackmann-Sala et al. 2014, Kwon et al. 2016, Sackmann Sala et al. 2017). The transcriptomic signature of LSC^med^ (*versus* other epithelial cells) identified them as a distinct epithelial cell entity, and CK4 was validated as a specific protein biomarker of LSC^med^ progenitor cells on prostate sections from various mouse models (Sackmann Sala et al. 2017). Within the past 12 months, several groups published single cell (sc) atlases of the adult mouse prostate cells based on RNA sequencing (scRNA-seq) data (Crowley et al. 2020, Guo et al. 2020, Joseph et al. 2020, Karthaus et al. 2020, Mevel et al. 2020). All studies identified one computerized cluster of non-secretory luminal progenitor cells enriched in luminal and stemness-related gene transcripts. The correspondence between the luminal progenitor cell clusters described in each scRNA-seq study was not investigated beyond the qualitative overlap of a few common markers. Furthermore, the presence of *Krt4* (encoding CK4) among the latter raised the question of the overlap between *in silico*-identified luminal progenitors (Crowley et al. 2020, Guo et al. 2020, Joseph et al. 2020, Karthaus et al. 2020, Mevel et al. 2020) and FACS-enriched LSC^med^ cells (Sackmann Sala et al. 2017). To address these questions, we used a unified bioinformatics pipeline to re-analyze the published prostate scRNA-seq datasets and perform various pan-transcriptomic comparisons. The data presented here demonstrate that i) the mouse prostate luminal progenitor cell clusters identified in the different scRNA-seq studies largely overlap and can be defined by a common 15-gene signature, ii) mouse LSC^med^ cells match both mouse and human luminal progenitors identified by scRNA-seq analysis.

### Experimental procedures

#### scRNA-seq data retrieval

Data re-analyzed as part of this study was retrieved from public databases with accession numbers GSE145861, GSE145865, GSE146811, GSE150692, GSE151944 (GEO, Gene Expression Omnibus) and OEP000825 (NODE, The National Omics Data Encyclopedia) (**Table 1**). We only retained samples taken from intact (i.e. non-castrated) mice. In particular, data retrieved from GEO repository GSE151944 were available as MULTI-seq sample barcodes; these were demultiplexed using the *MULTIseqDemux* function implemented in *Seurat* (Stuart et al. 2019). For datasets GSE145861 and GSE145865 (Joseph et al. 2020), the data from the prostate and urethral regions were aggregated during the analysis step. Finally, data retrieved from the OEP000825 repository were raw fastq files which were processed (read alignment, generation of feature-barcode matrices) with Cell Ranger (10x Genomics) prior to any data analysis step.

**Table 1.**

| **Repository** | **Accession** | **Data format** | **Cell count** | **Reference** | **DOI** |
| --- | --- | --- | --- | --- | --- |
| GEO (Gene Expression Omnibus) | GSE145861<br>GSE145865 | Count matrices in h5 format for each sample | 90345 | Joseph et al.<br>(The Prostate, 2020) | 10.1002/pros.24020 |
| GEO (Gene Expression Omnibus) | GSE146811 | Pooled Count matrices in h5 format | 13688 | Karthaus et al.<br>(Science, 2020) | 10.1126/science.aay0267 |
| NODE (National Omics Data Encyclopedia) | OEP000825 | Raw fastq files | 34444 | Guo et al.<br>(Nat Genetics, 2020) | 10.1038/s41588-020-0642-1 |
| GEO (Gene Expression Omnibus) | GSE150692 | Raw counts matrices in tsv format for each sample | 5288 | Crowley et al.<br>(eLife, 2020) | 10.7554/eLife.59465 |
| GEO (Gene Expression Omnibus) | GSE151944 | MULTI-seq outputs as raw count matrices per sample | 4624 | Mevel et al.<br>(eLife, 2020) | 10.7554/eLife.60225 |

#### Data analysis

All datasets were processed using the same analytical pipeline. Low quality cells were filtered-out by consecutively filtering each sample individually (i.e. prior to any aggregation step) based on unique molecular identifiers (UMI) counts, percentage mitochondrial content and number of genes (in that precise order). No filter was applied on the percentage of ribosomal gene content. Filtering was performed as previously described (Henry et al. 2018, Joseph et al. 2020): filter thresholds were chosen dynamically for samples based on the distribution of each parameter, with code adapted from sc-TissueMapper (v2.0.0) (Henry and Strand 2020). Upper and lower filters were applied on UMIs (the lower bound of the UMI filter was strictly set to 200 whilst the *RenyiEntropy* thresholding technique was applied to determine the upper bound after binning the data), while the percentage mitochondrial content had only an upper filter (abnormally high percentages of mitochondrial content were determined using the *Triangle* filter on binned data) and feature number had only lower filters (determined using the *MinErrorI* filter on binned data). *RenyiEntropy, Triangular*, and *MinErrorI* thresholding were applied using functions from the *autothresholdr* (v1.3.9) R package. For a given dataset, if multiple samples were available, samples were aggregated by normalizing with the *sctransform* (version 0.3.2) method and using *Seurat*’s reciprocal PCA method. Furthermore, cells displaying high stress signatures associated with the tissue dissociation experimental step were removed as described (Henry et al. 2018): aggregated cells were scored for stress using *Seurat*’s *AddModuleScore* method and a mouse geneset enriched for stressed cells earlier described (van den Brink et al. 2017). Finally, principal component analysis (PCA) was performed on the data and graph-based clustering was performed using the principal components representing 90% of the associated cumulative variance.

#### LSC^med^ score calculation

LSC^med^-specific genes (*n* = 111 genes, assayed in WT mice) were retrieved from Sackmann Sala *et al*. (2017). The LSC^med^ similarity score was based on the calculation of the average mRNA levels of the 111 signature genes for each single cell (to which is subtracted the averaged mRNA levels of 50 randomly chosen control genes) using *Seurat*’s *AddModuleScore* method. For calculation in the human prostate dataset, mouse stable gene IDs were matched to human stable gene IDs, retaining one-to-one ortholog matches. The mouse-human orthology table was generated with Ensembl BioMart.

#### Marker gene identification

Differentially expressed genes for the identified cell subpopulations were determined using Wilcoxon rank sum tests on genes present in at least 20% of cells in the population of interest, only retaining positive gene markers. Testing was limited to genes which showed, on average, at least 0.2-fold difference (on a log-scale) between the different groups. Finally, genes displaying an adjusted *p*-value inferior to 5% (*P*_adj_ < 0.05) were retained.

#### Data access

The scRNA-seq datasets re-analyzed in the scope of this project were stored as processed R objects in .rds format and are available from the corresponding author on reasonable request.

## Results

To investigate the overlap between the luminal progenitor clusters described in published single cell studies (Crowley et al. 2020, Guo et al. 2020, Joseph et al. 2020, Karthaus et al. 2020, Mevel et al. 2020), we reanalyzed their publicly available scRNA-seq transcriptomic data (**Table 1**) to identify the top biomarkers of luminal progenitors in each individual study and compare their expression level in each cell population across studies. Importantly, we applied the same analytical pipeline on count data retrieved for each of the studies, allowing notably to identify and remove stressed cells from the analyses (see **Experimental procedures**, *Data analysis*), enabling comparisons between the different datasets. Starting from 4,624, 13,688, 5,288, 34,444 and 90,345 single cells for each study (**Table 1**), we retained 1,213, 5,158, 2,362, 19,503 and 45,432 single cells after quality filtering, respectively (see Legend to **Figure 1A**), with a median of 2,205 genes assayed per cell.

**Figure 1.**
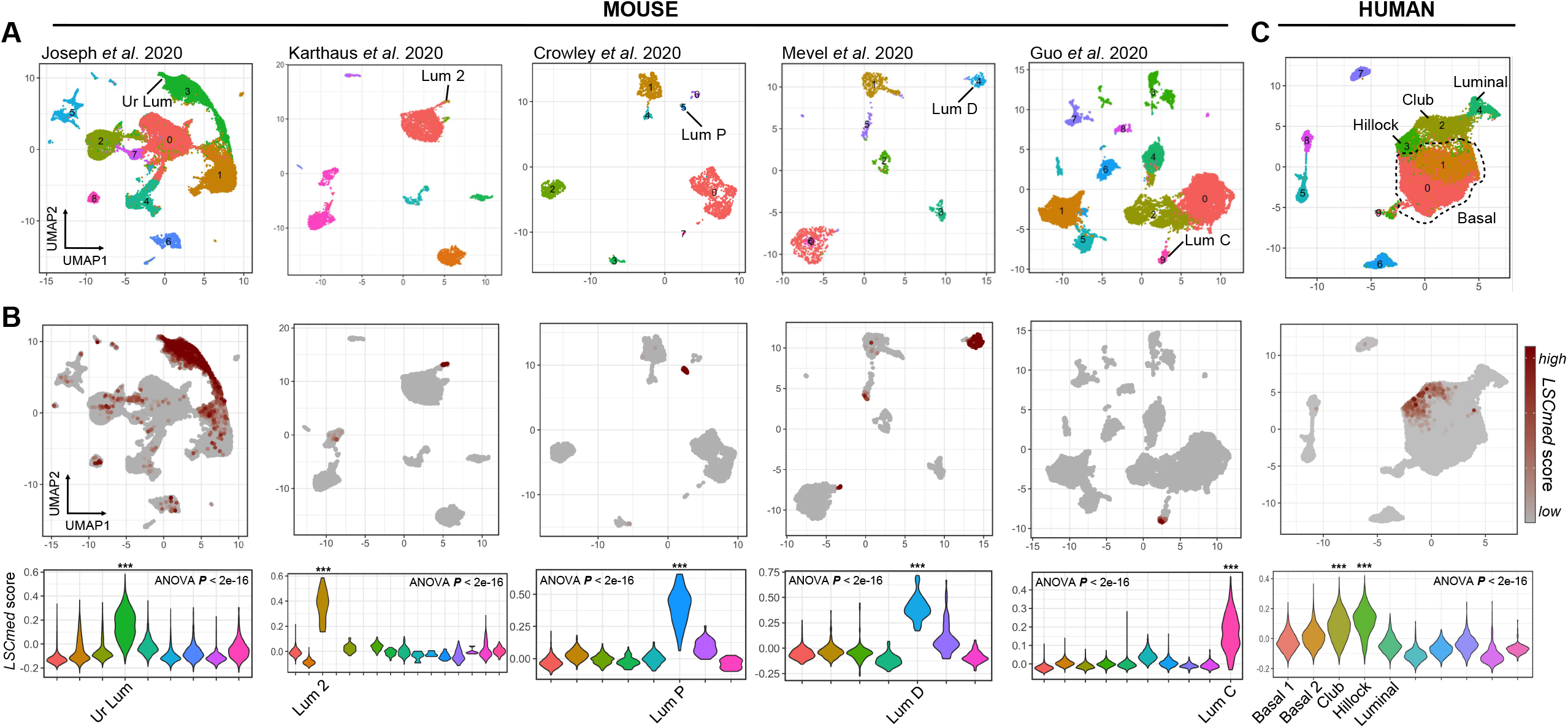
Transcriptomic similarity between FACS-enriched WT mouse LSC^med^ cells and luminal progenitor cell clusters identified by scRNA-seq analyses of mouse (A,B) and human (C) healthy prostates. **(A)** UMAP projections based on linear dimensionality reduction by principal component analysis (PCA) for, from left to right, 45,432 (Joseph et al. 2020), 5,158 (Karthaus et al. 2020), 2,362 (Crowley et al. 2020), 1,213 (Mevel et al. 2020) and 19,503 (Guo et al. 2020) single-cell transcriptomes. **(B)** In each study, a single subpopulation matched LSC^med^-like cells (Sackmann Sala et al. 2017), as shown by high calculated LSC^med^ gene signature scores. The violin plots show the calculated LSC^med^ scores per cluster and per study (^***^, Tukey multiple comparisons of means *P*_*adj*_ < 0.001 for all pairwise comparisons performed). **(C)** The same analysis was performed for human prostate single-cell transcriptomes (Joseph et al. 2020).

Subsequently, clustering was followed by dimensional reduction for visualization using UMAP (Uniform Manifold Approximation and Projection) plots, which depicts cell populations with distinct transcriptional signatures (**Figure 1A**). For each dataset, we identified between 7 and 14 *in silico* computed clusters expressing different phenotypic markers (epithelial, immune and stromal cells) as described (Crowley et al. 2020, Guo et al. 2020, Joseph et al. 2020, Karthaus et al. 2020, Mevel et al. 2020), which were used to match previously labeled sub-populations of interest to our re-analyzed datasets (**Figure 1A**). In particular, we were able to recapitulate the Lum D (Mevel et al. 2020), Lum 2 (Karthaus et al. 2020), Lum P (Crowley et al. 2020), Lum C (Guo et al. 2020) and Ur Lum (Joseph et al. 2020) subsets of non-secretory luminal progenitor cells as clear clusters. Marker gene identification for these subpopulations (**Table S1**) allowed to identify 21 different genes common to at least 4 studies, of which 15 were common to all studies (**Figure 2**). These included *Krt4, Psca, Clu, Wfdc2, Cyp2f2, Tspan8, Gsta4 and Tacstd2*. Of note, the latter has also been used as a surface protein biomarker (Trop2) to enrich luminal progenitors by cell sorting (Crowell et al. 2019, Crowley et al. 2020, Guo et al. 2020). This analysis demonstrates that the various luminal progenitor clusters identified by scRNA-seq exhibit a high degree of similarity and can be defined by a common transcriptomic signature.

**Figure 2.**
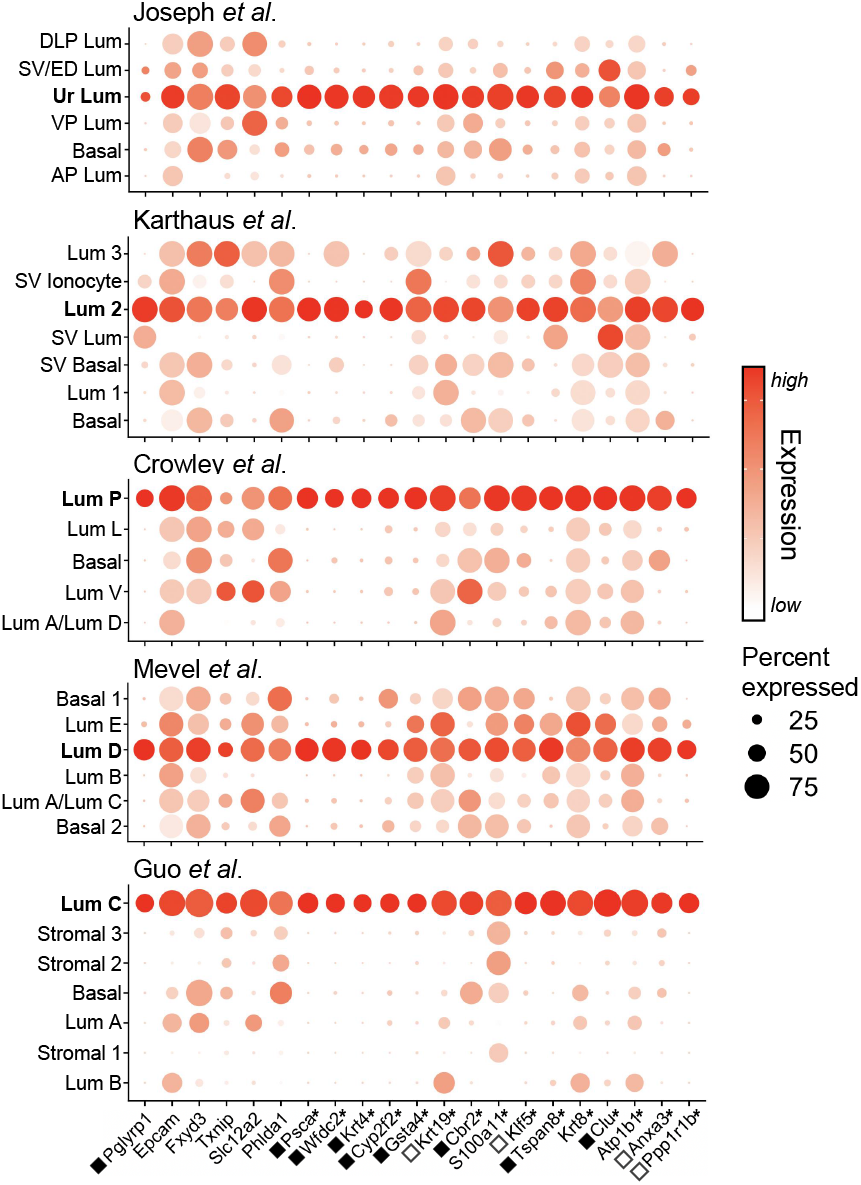
Common markers of luminal progenitors identified by scRNA-seq study of WT mouse prostate. Dot plot representation of marker genes for LSC^med^-like cell subpopulations found in the five scRNA-seq studies discussed in the text (as indicated), showing their relative expression across all detected clusters. Each dot depicts both detection rate and average gene expression in detected cells for a gene in a cluster. Darker red colors indicate higher average gene expression, and a larger dot diameter indicates that the gene was detected in greater proportion of cells from the cluster. Stars (*) depict LSC^med^-like cell marker genes detected in all datasets. Squares indicate marker genes that were significantly (black) or nearly significantly (white) identified in the LSC^med^ cell signature (Sackmann Sala et al. 2017). The other genes were all expressed in LSC^med^ cells at similar levels as in basal and/or luminal cells, therefore they were not identified as LSC^med^ cell markers.

All above-mentioned genes are part of the LSC^med^ signature (Sackmann Sala et al. 2017). To assess the actual overlap of LSC^med^ cells with luminal progenitors identified by scRNA-seq, we sought for the enrichment of the LSC^med^ cell signature (111 genes) in the various prostate cell populations identified *in silico* (see **Experimental procedures**, *LSCmed score calculation*). As expected, significant enrichment of the LSC^med^ cell signature was observed in each scRNA-seq dataset for the sole population identified as luminal progenitors (**Figure 1A**,**B**). Altogether, this indicates that LSC^med^ progenitor cells enriched by cell sorting (Sackmann-Sala et al. 2014, Kwon et al. 2016, Sackmann Sala et al. 2017) largely overlap with luminal progenitor cell clusters identified by scRNA-seq.

Single cell analyses of the human prostate have also been recently reported (Henry et al. 2018, Crowley et al. 2020, Joseph et al. 2020, Karthaus et al. 2020). The original report by the group of D. Strand identified two closely-related clusters of luminal progenitor-like cells showing transcriptomic similarities with Club and Hillock progenitor cells described in the lung (Henry et al. 2018) (**Figure 1C**). Prostate Club/Hillock cells share typical markers with mouse luminal progenitors including as *KRT4, TACSTD2* and *PSCA*. As observed above for mouse luminal progenitors (**Figure 1A,B**), the LSC^med^ cell signature was significantly enriched in Club and Hillock progenitors, but not in other cells of the human prostate (**Figure 1C**).

Together, the full transcriptomic profile comparisons reported above provide unbiased evidence that mouse prostate LSC^med^ luminal progenitors enriched by cell sorting correspond to luminal progenitor clusters identified in mouse and human prostates by scRNA-seq.

## Discussion

Luminal progenitors have recently emerged as key players of prostate pathogenesis including inflammation (Wang et al. 2015), benign prostate hyperplasia (Crowell et al. 2019, Joseph et al. 2020) and prostate cancer (Korsten et al. 2009, Wang et al. 2014, Sackmann Sala et al. 2017, Guo et al. 2020). These findings have generated a particular interest for this particular cell subset (Joseph et al. 2021). However, one recurrent issue in the stem cell field is to evaluate to which extent cell populations identified in different studies, often by different experimental approaches, actually correspond to equivalent cell entities. While the probable overlap of *in silico* computed clusters of prostate luminal progenitors in the various scRNA-seq studies was suggested (Crowley et al. 2020), to our knowledge, no *bona fide* transcriptomic analysis covering all scRNA-seq data available was provided to support this hypothesis. Furthermore, despite of the fact that CK4 was stressed as a biomarker of the luminal progenitor cluster in at least two studies (Guo et al. 2020, Mevel et al. 2020), the potential similarity with LSC^med^ cells was ignored in all but one scRNA-seq report (Joseph et al. 2020). The present study involving pan-transcriptomic comparative analysis of human and mouse luminal progenitor clusters identified *in silico versus* enriched *ex vivo* by cell sorting definitely confirms their molecular equivalence which is materialized by the common 15-gene signature. We and others demonstrated that mouse prostate LSC^med^ luminal progenitors can initiate tumors in reconstitution assays and are castration-tolerant in both normal and pathological states (Kwon et al. 2016, Sackmann Sala et al. 2017, Karthaus et al. 2020), suggesting they may contribute to resistance to treatments targeting androgen signaling. In keeping with this, some scRNA-seq studies highlighted the transcriptional plasticity of epithelial luminal cells upon castration (Karthaus et al. 2020, Mevel et al. 2020). Bridging such *in silico* transcriptomic information with functional characterization of FACS-enriched luminal progenitor cells knowing they apply to the same cell entity should speed up our understanding of their biology in health and disease.

## Acknowledgements

The authors wish to thank la Ligue contre le cancer (RS20/75-93 and RS21 /75-35), FONCER contre le cancer, Inserm and the University Paris Descartes. M.B. is supported by a fellowship from the Ministry of Research, C.D. by research/mobility fellowships from Inserm, the Association Française d’Urologie and Assistance Publique Hôpitaux de Paris (APHP).

